# Expression of mutant TIE2 p.L914F during mouse development causes embryonic lethality and defects in vascular remodeling

**DOI:** 10.1101/2025.07.24.666614

**Authors:** Lindsay J Bischoff, Sandra Schrenk, Kara Soroko, Elisa Boscolo

## Abstract

**Background:** Sporadic venous malformation (VM) is associated with the hyperactivating p.L914F mutation in TIE2, a receptor tyrosine kinase essential for vascular development. This mutation is not found in hereditary VM, suggesting incompatibility with life when expressed during early vascular development. Therefore, we utilized a novel genetic mouse model that expresses TIE2 p.L914F to determine the phenotypical effects of this mutation during development.

**Results:** *B6-Tg(Rosa26-TIE2^L914F^)^EBos^* (*TIE2^L914F^*) mice were generated and then validated for the presence of the transgene. The constitutive endothelial-specific *Tie2-Cre* line was used to activate expression of the mutant gene during early embryonic development. *Tie2-Cre;TIE2^L914F^*embryos experienced lethality at approximately embryonic day (E)9.5. 3-dimensional imaging of embryos and yolk sacs revealed impaired vascular remodeling in mutant animals, resulting in malformed vasculature with disorganized, dilated, and non-functional blood vessels. The abnormal vascular phenotype was not associated with total loss of erythroid cells or increased cell proliferation.

**Conclusions:** The *TIE2^L914F^* mice used in this study represent a novel genetic model of TIE2 p.L914F-driven vascular disease. This study provides the first experimental evidence that this mutation is incompatible with early prenatal development due to its deleterious effects on the vasculature, illustrating the vital role of TIE2 signaling during vessel development and remodeling.

## Introduction

Vascular malformations encompass a group of disorders thought to arise from errors in physiological vascular development^1^. Venous malformation (VM) is the most common type of vascular malformation, with an estimated incidence of 1-2 per 10,000 births^2,3^. VM lesions consist of dilated slow flow veins which can cause significant morbidity, including disfigurement, chronic pain, and coagulopathy^4–7^. VM lesions can be congenital or make their appearance during childhood, but never regress without intervention^4^. Activating mutations in the *TEK* gene, which encodes for the TIE2 receptor tyrosine kinase, have been identified in the majority of VM cases^8–11^. Interestingly, TIE2 causative mutations have been found to be both germline (causing rare hereditary forms of disease) and somatic (causing sporadic forms)^12^. In the sporadic cases, the TIE2 mutation has been identified specifically in lesional endothelial cells (ECs)^13^, implicating these cells as the major driver of VM development.

Most VM-associated TIE2 mutations are activating (causing aberrant ligand-independent autophosphorylation of the receptor^8,11,14–16)^, however there are differences in the level of receptor activation and patterns of downstream signaling amongst the different mutations. The mutations that have been identified in the germline (such as p.R849W) tend to have lower receptor auto-phosphorylation and weaker downstream signaling than those that are only found somatically (such as p.L914F)^16^. These differences in the magnitude of changes downstream of the mutant receptor have been hypothesized to explain the differences in germline vs somatic occurrence of these mutations. While hereditary mutations are sometimes found in sporadic VM, the most common somatic mutations never appear in the germline, perhaps due to stronger signaling activation being incompatible with life. The most prevalent somatic mutation is p.L914F, which has been identified in up to 80% of TIE2-related somatic cases, but has never been found in hereditary forms of the disease^9^. This has led to the hypothesis that TIE2 p.L914F mutation would be embryonically lethal if expressed in the germline or early during embryonic development, but will be tolerated in the mosaic or somatic condition, as proposed for other developmental and sporadic disorders^17,18^. For other somatic activating mutations found in vascular anomalies, such as GNAQ^19,20^ and PIK3CA^21^, genetic mouse models have revealed that early developmental expression causes embryonic lethality and vascular defects. However, due to the lack of a genetic animal model expressing mutant TIE2, the above-mentioned hypothesis has never been empirically tested.

TIE2 expression is enriched in ECs^22^ and has critical roles in vascular development^23^, stability and quiescence^24^, remodeling^25^, and angiogenesis^26^. TIE2 activation is tightly balanced in the vasculature to facilitate these functions. TIE2 phosphorylation has been shown in both quiescent and angiogenic vessels^27^, indicating its dual roles in stable and active ECs. TIE2 signaling activation primarily occurs as a consequence of binding to its ligand angiopoietin-1 (ANGPT-1), while angiopoietin-2 (ANGPT-2) can act as both antagonist or activator of the receptor, depending on the environmental and cellular context^28,29^. Deletion of TIE2^30,31^ or ANGPT-1^32^, or overexpression of ANGPT-2^33^, during embryogenesis results in lethality due to severe vascular defects, including failure of vascular remodeling (summarized in ^23^). However, the effects of TIE2 hyperactivation in the developing vasculature have not yet been investigated.

Therefore, in the present study, we have generated a new genetic mouse model that conditionally expresses TIE2^*L914F*^. In order to investigate the effects of this mutation during early development in ECs, we crossed the *TIE2^L914F^* mice to a constitutive endothelial-specific *Tie2-Cre* driver and examined embryonic survival and vascular phenotypes.

## Results

### Generation of a genetic mouse model with conditional expression of mutant TIE2 p.L914F

In previous studies, injection of ECs expressing the TIE2 p.L914F mutation in mouse xenografts established that this mutation is sufficient to confer them the ability to form aberrant vascular structures reminiscent of VM^13,34,35^. Here, we aimed to investigate the effects of the TIE2 hyperactivating mutation during embryonic development by generating a genetic murine model for conditional expression of the TIE2 p.L914F mutation. To do so, we constructed a *Rosa26*-targeting vector containing the coding sequence of the human *TEK* gene with the p.L914F (c.2740C>T) mutation. The expression of the *TEK* transgene is controlled by a Neo-STOP cassette, flanked by LoxP sites (floxed) (**Figure 1A**), which prevents transgene expression until Cre recombination occurs. Homology-mediated recombination was used to achieve targeted knock-in of the transgenic vector into the *Rosa26* locus of C57BL/6 mice (*B6-Tg(Rosa26-TIE2^L914F^)^EBos^ –* thereafter called *TIE2^L914F^*).

**Figure 1:**
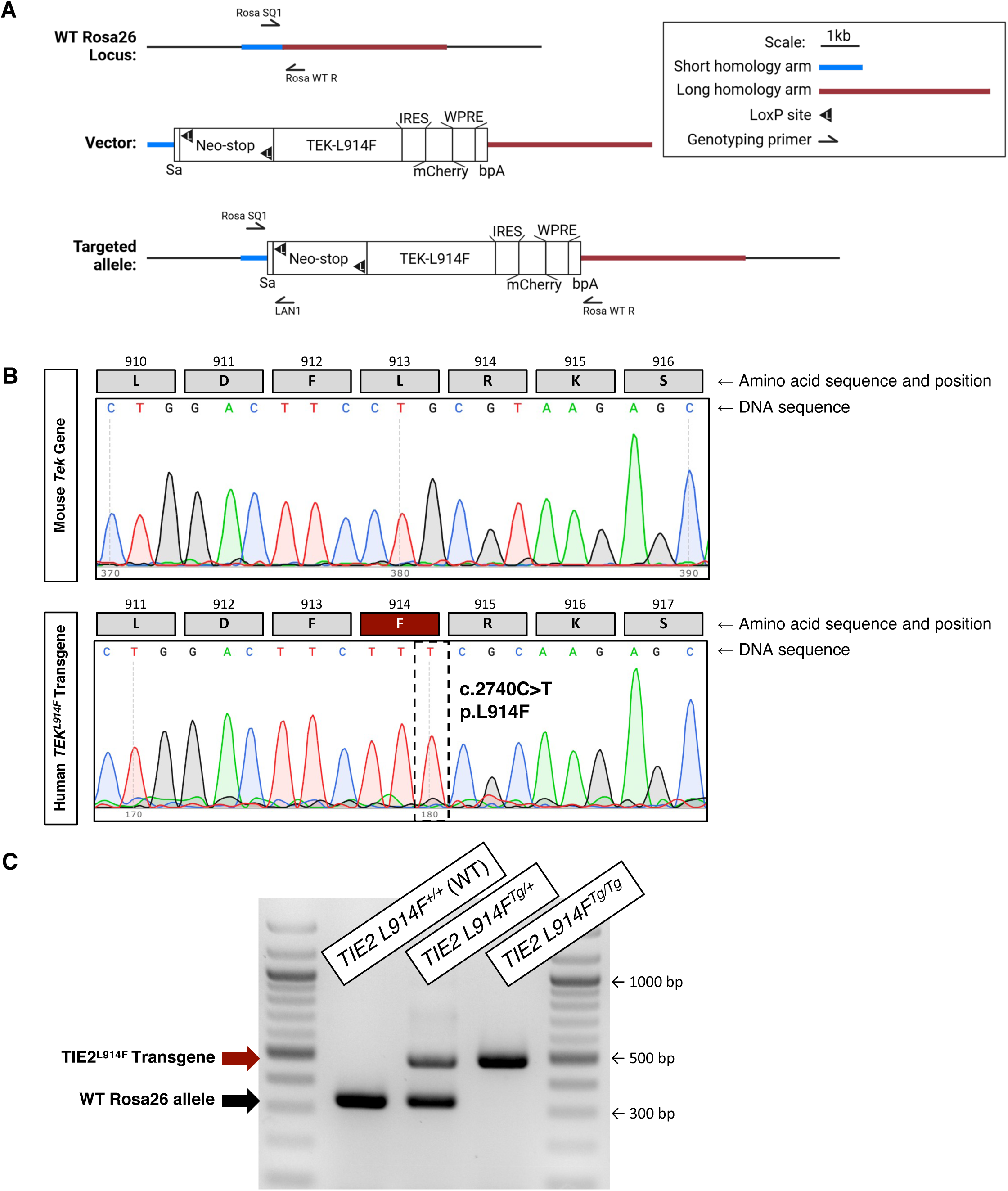
Generation of a *Rosa26-floxed stop-TIE2^L914F^*(*TIE2^L914F^*) mouse. (A) Schematic of the transgenic targeting vector and integration into the *Rosa26* mouse locus. Sa=splice acceptor, IRES=internal ribosome entry site, WPRE=woodchuck hepatitis virus posttranscriptional regulatory element, bpA=bovine growth hormone polyadenylation signal. (B) Chromatograms from Sanger sequencing of Exon 17 of the endogenous mouse *Tek* gene and human *TEK* transgene from a *TIE2^L914F^* mouse. The corresponding amino acid and position of each in-frame codon is shown above. Due to differences between the mouse and human orthologs, leucine 914 in human corresponds to leucine 913 in mice. The black dashed box indicates the single nucleotide substitution that results in the p.L914F mutation in the human transgene. (C) Representative genotyping results from mice harboring zero, one, or two copies of the TIE2^*L914F*^ transgene (Tg). (+) represents the WT *Rosa26* allele. Using the three-primer protocol, the WT *Rosa26* locus and the integrated transgene can be simultaneously amplified, resulting in two bands at approximately 330bp and 500bp, respectively. bp = base pairs.

To confirm the presence of the p.L914F (c.2740C>T) mutation, Exon 17 of *TEK* was specifically amplified from both the endogenous mouse locus and the human transgene of *TIE2^L914F^*mice. Sanger sequencing revealed the p.L914F mutation in the transgene, which was absent in the wild-type (WT) mouse *Tek* allele (**Figure 1B**). TIE2 is highly conserved between species, with the human and murine genes sharing 88 and 92 percent sequence homology of the coding DNA and amino acid sequences, respectively (Ensembl).

We then designed a 3-primer genotyping protocol utilizing one common forward primer (Rosa SQ1), one Rosa-specific reverse primer (Rosa WT R), and one transgene-specific reverse primer (LAN1). Utilizing this protocol, we can simultaneously amplify the WT Rosa26 integration site, as well as the integrated transgene. The products of each amplification reaction differ in size, so we can distinguish between mice harboring zero, one, or two copies of the integrated allele (**Figure 1C**). *TIE2^L914F^* mice are overtly phenotypically normal and indistinguishable from WT C57/BL6 mice. Over 6 generations of colony breeding, we have observed no obvious deleterious effects on mouse fertility, health, or survival. In total, these data demonstrate the successful generation of a novel genetic mouse model containing a Cre-inducible *TIE2^L914F^* transgene.

### Endothelial-specific expression of mutant TIE2 p.L914F during embryonic development results in mid-gestation lethality

In humans, the TIE2 p.L914F mutation has been identified in VM patients with somatic or mosaic expression pattern. We hypothesized that constitutive expression of mutant TIE2 p.L914F starting during early embryonic vascular development would be incompatible with life. To test this hypothesis, we crossed *TIE2^L914F^* mice to the *Tg(Tek-cre*)*^1Ywa^*line (*Tie2-Cre*), which has a *Tek* promoter region linked to Cre recombinase for expression specifically in cells of endothelial lineage, beginning at approximately embryonic day 7.5 (E7.5)^36^. Upon mating *Tie2-Cre^Tg/+^* males to *TIE2^L914F^ ^Tg/+^*females (**Figure 2A**), we observed no *Tie2-Cre;TIE2^L914F^*pups, compared to the Mendelian expected 25% of the offspring (**Table 1**). These results suggest that early embryonic expression of mutant TIE2 p.L914F in ECs is embryonically lethal.

**Figure 2:**
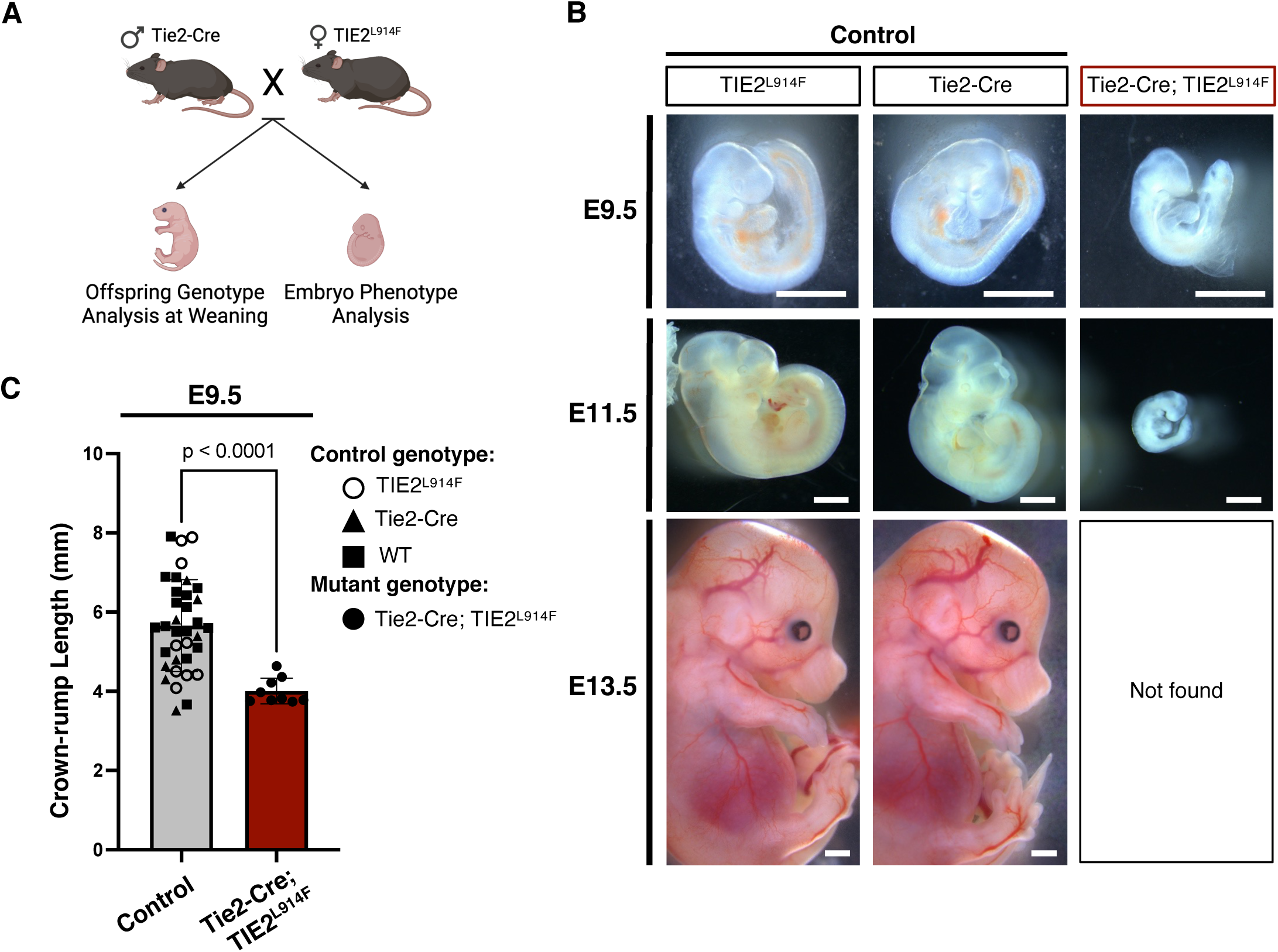
Constitutive and endothelial-specific TIE2^L914F^ expression causes embryonic lethality. (A) Schematic of the *Tie2-Cre, TIE2^L914F^* breeding scheme and downstream analyses. (B) Representative gross images of mouse embryos collected at the indicated timepoints. At E13.5, no *Tie2-Cre; TIE2^L914F^* embryos were found. Remnants of yolk sacs were used to confirm genotypes of empty implantation sites. n=35 controls and n=9 mutants at E9.5, n=12 controls and n=9 mutants at E11.5, n=11 controls and n=5 mutants at E13.5. Scale bars=2.5mm. (C) Length measurements from the crown to the rump of E9.5 embryos. Control specimens include embryos of each genotype, indicated in the figure legend. WT=wild-type (transgene negative). n=35 controls and n=9 mutants. Mean±SD. Welch’s t-test.

**Table 1:** Expected and observed genotypes of offspring from *Tie2-Cre* x *TIE2^L914F^* breeding. Tie2-Cre^Tg/+^ mice were bred with TIE2 L914F^Tg/+^ mice. Hemizygous x hemizygous crosses are expected to yield 25% of each possible genotype in offspring. Genotyping was performed on live offspring at weaning (postnatal day 21). The actual percentage and total number of offspring observed is listed in the rightmost column. A chi-square goodness-of-fit test was performed to calculate the p-value of the difference between the expected and observed genotypes (below table). Tg indicates one copy of the transgene. (+) indicates one copy of the wild-type allele. WT=wild-type (negative for both transgenes).

| Tie2-Cre<sup>Tg/+</sup> x TIE2 L914F<sup>Tg/+</sup> Offspring Genotype |  |  |  |
| --- | --- | --- | --- |
| Group | Genotype | Expected | Actual<br>(total = 42) |
| WT | Tie2-Cre<sup>+/+</sup> ; TIE2 L914F<sup>+/+</sup> | 25% | 26% (n=11) |
| Tie2-Cre | Tie2-Cre<sup>Tg/+</sup> ; TIE2 L914F<sup>+/+</sup> | 25% | 42% (n=18) |
| TIE2<sup>L914F</sup> | Tie2-Cre<sup>+/+</sup> ; TIE2 L914F<sup>Tg/+</sup> | 25% | 31% (n=13) |
| Tie2-Cre;TIE2<sup>L914F</sup> | Tie2-Cre<sup>Tg/+</sup> ; TIE2 L914F<sup>Tg/+</sup> | 25% | 0% (n=0) |

Chi-square value = 16.48, p = .0009

To determine the embryonic stage at which lethality occurred, we performed timed mating of *Tie2-Cre^Tg/+^* males and *TIE2 ^L^*^914^*^F^ ^Tg/+^* females. Embryos were collected at E9.5, 11.5, and 13.5 (**Figure 2B**). At E9.5, control and mutant embryos were grossly similar, although the mutants appeared to have a growth delay and were more pale in comparison to controls, suggesting an absence of blood. Analysis of the crown-to-rump length at E9.5 showed that mutant embryos were significantly smaller in size (**Figure 2C**). At this time point, mutant embryos were likely still viable, as we observed heartbeats in both control and mutant specimens (**Videos 1** and **2**). However, by E11.5, all mutant embryos failed to develop further and were deceased. At E13.5, no *Tie2-Cre;TIE2^L914F^*embryos were found and had undergone resorption, resulting in empty implantation sites. Together, these results suggest that TIE2^*L914F*^-mutant embryos survive until approximately E9.5 but fail to develop beyond this timepoint.

### *Tie2-Cre;TIE2^L914F^* embryos exhibit aberrant dilation and remodeling of the vasculature

To determine whether mutation-induced changes to the vasculature were responsible for the observed embryonic lethality, we performed whole-mount immunostaining and confocal imaging of embryos at E9.5. CD31 endothelial-specific labeling of the vasculature (**Figure 3A**) revealed several defects in the remodeling of the vascular plexus and aberrant vessel size in *Tie2-Cre;TIE2^L914F^* embryos when compared to controls. We utilized Nikon software to construct depth-coded, volume views of whole embryos, as well as specific structures (**Figure 3B-D**). We first noted that the cephalic plexus of the cranial region was massively disorganized in *Tie2-Cre;TIE2^L914F^* embryos (**Figure 3C**). During normal embryonic development, the cephalic plexus arises from CD31-expressing clusters of cells within the cephalic mesoderm that establish a primitive plexus, which ultimately remodels to form a hierarchical honeycomb-like pattern of vessels^37^. Whereas these vessels formed a distinctive network in the control animals, they were disorganized and dilated in the mutants, indicating a defect in remodeling of the primitive cephalic plexus.

**Figure 3:**
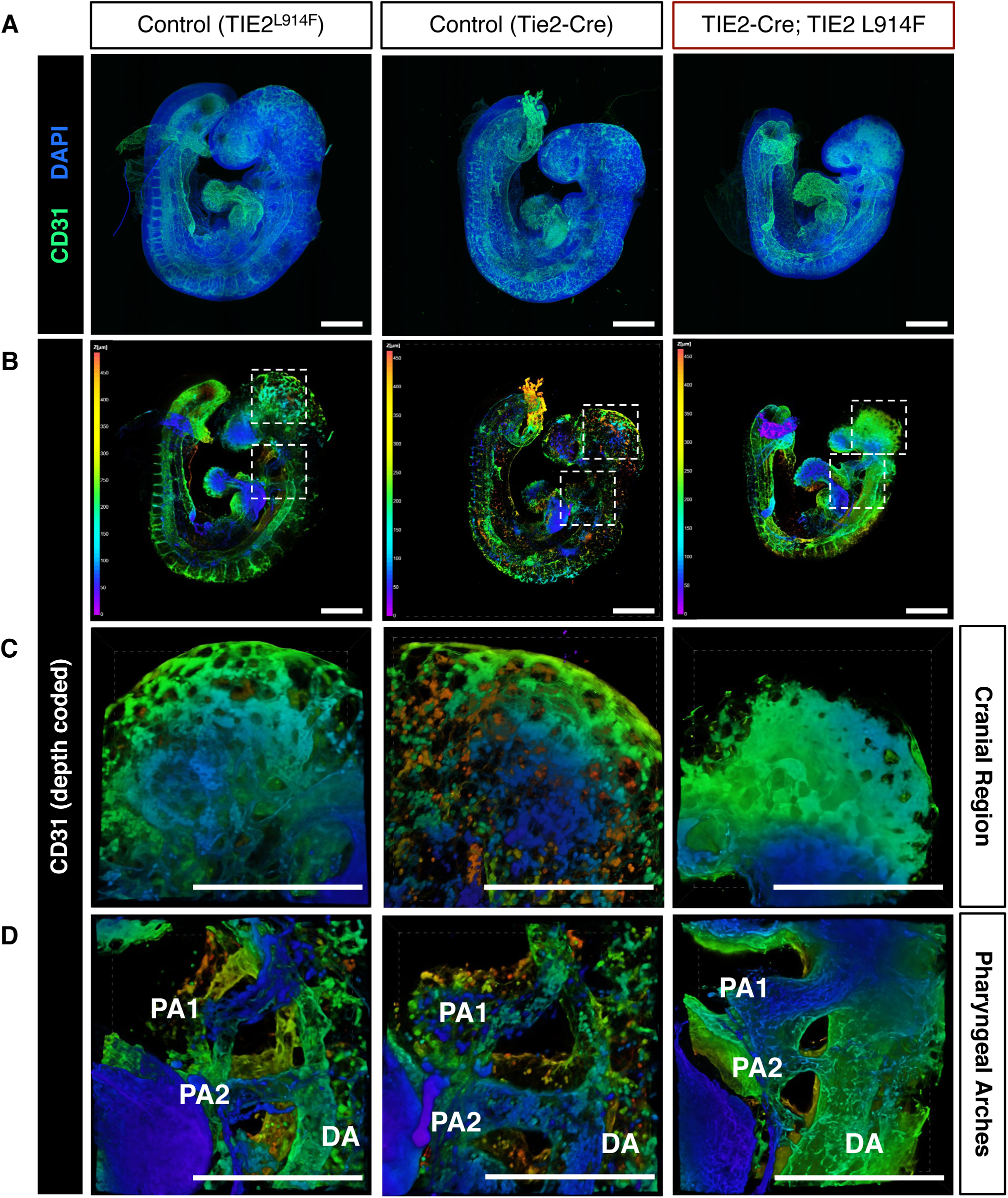
TIE2^L914F^ expression causes aberrant development of the vasculature in E9.5 embryos. (A) Representative maximum intensity projections of tile-scan z-stack confocal images. Whole-mount E9.5 embryos of the indicated genotypes were immunostained for the vasculature (CD31, green) and nuclei counterstained with DAPI (blue). (B) Nikon Elements software was used to construct volume views of the images in (A), with CD31 signal depth-coded according to its location on the z-axis. Depth-coding color scale shown to the left in images. White dashed boxes show location of zoomed images in (C-D). High resolution magnified images of anatomic locations from tile-scan. Shown are the cranial region (C) and pharyngeal arches (D). PA1 or 2=pharyngeal arch artery 1 or 2, DA=dorsal aorta. Additional embryonic locations (heart and somites) are shown in Supplemental Figure 2. All scale bars are 500µm. Representative of n=9 controls and n=7 mutants.

Furthermore, we observed the presence of the first and second pharyngeal arch arteries in the control embryos (**Figure 3D**), which form at 9-9.5 days of gestation and eventually regress and remodel into other components of the arterial system by E10-10.5^38^. In the *Tie2-Cre;TIE2^L914F^*embryos, we found there was massive dilation of the first and second pharyngeal arch arteries, accompanied by dilation of the dorsal aorta. The dilation of these structures can be clearly observed via digital sectioning of volume reconstructions of the embryo wholemounts (**Supplementary Figure 1**). We also examined the intersomitic vessels (ISVs), which sprout from the dorsal aorta and cardinal veins and migrate dorsally between the somites to form the vertebral artery, dorsal longitudinal anastomotical vessel, and the perineural vascular plexus^37^. Here, we observed normal sprouting of the ISVs in *Tie2-Cre;TIE2^L914F^* embryos (**Supplementary Figure 2**).

As formation and remodeling of the embryonic vasculature is influenced by flow-mediated signaling from blood circulation^39–42^, we also examined the vasculature of the heart. At E9.5, the heart is apparent as a looped heart tube consisting of the primitive atria and ventricles, with the inflow and outflow tracts connecting the heart to the venous and atrial circulation, respectively^43^. Morphologically, these structures appeared similar in the controls and *Tie2-Cre;TIE2^L914F^*embryos indicating that, to this point, cardiac development is unaffected by the TIE2 p.L914F mutation (**Supplementary Figure 2**). Preservation of heart function was also supported by our observations of normal cardiac contractions in isolated control and mutant embryos (**Videos 1** and **2**).

Together, these data demonstrate that developmental expression of TIE2 p.L914F in ECs causes aberrant formation and dilation of the embryonic vasculature of the cranial and pharyngeal arches regions, by E9.5.

### *Tie2-Cre;TIE2^L914F^* embryos show massive defects of the yolk sac vasculature

As defects in the embryonic vasculature are often also associated with defects in the yolk sac vasculature, we next examined the yolk sacs of control and mutant embryos. The vasculature of the yolk sac originates at E7.5 as collections of blood islands, which consist of blood and EC precursors^44^. Angioblasts from the blood islands then migrate to form a simple capillary plexus around E8.5^45,46^. From E8.5 to E9.5, this plexus undergoes rapid angiogenic remodeling to form the hierarchical network of large and small vessels that are necessary to support the circulation of the growing embryo^39,40^. Proper establishment of circulation is critical to ensure viability of the organism, therefore defects in the yolk sac vascular remodeling are highly associated with mortality at this timepoint.

Therefore, we analyzed the morphology and vasculature of control and mutant yolk sacs at E9.5. The yolk sacs of *Tie2-Cre;TIE2^L914F^* embryos were visibly distinct from those of controls, appearing abnormally wrinkled and deflated (**Figure 4A, top**). Closer examination of the yolk sacs revealed the presence of large, blood-filled vitelline vessels (arrows) in yolk sacs with control genotypes. In contrast, the mutant yolk sacs appeared to lack these networks of large vessels and instead showed areas of pooling blood (arrowheads) (**Figure 4A, bottom**). The clear presence of blood in the yolk sacs contrasts to the relative paleness of mutant embryos, perhaps indicating a defect in blood circulation from the yolk sac to the embryo. To examine the vasculature in greater detail, we performed immunostaining for CD31 of flat-mounted whole yolk sacs. With low-objective imaging, we observed substantial disorganization of the vasculature in *Tie2-Cre;TIE2^L914F^* yolk sacs (**Figure 4B, left**). At higher magnification, the formation of patterned vessels was completely abrogated in mutants, with ECs appearing to form sheets of cells (**Figure 4B, right**). Slice view through the z-plane of the whole mounts revealed two layers of cell sheets, with luminal space in between, and only few points of connection between the top and bottom layers. This was distinct from the sections of control yolk sacs, where the round lumens of patent vessels can be seen (**Figure 4B, bottom right**). This suggests that the disorganized vasculature of the *Tie2-Cre;TIE2^L914F^* yolk sacs acts as a continuous large vessel upon failure of the plexus to organize into the normal vascular tree.

**Figure 4:**
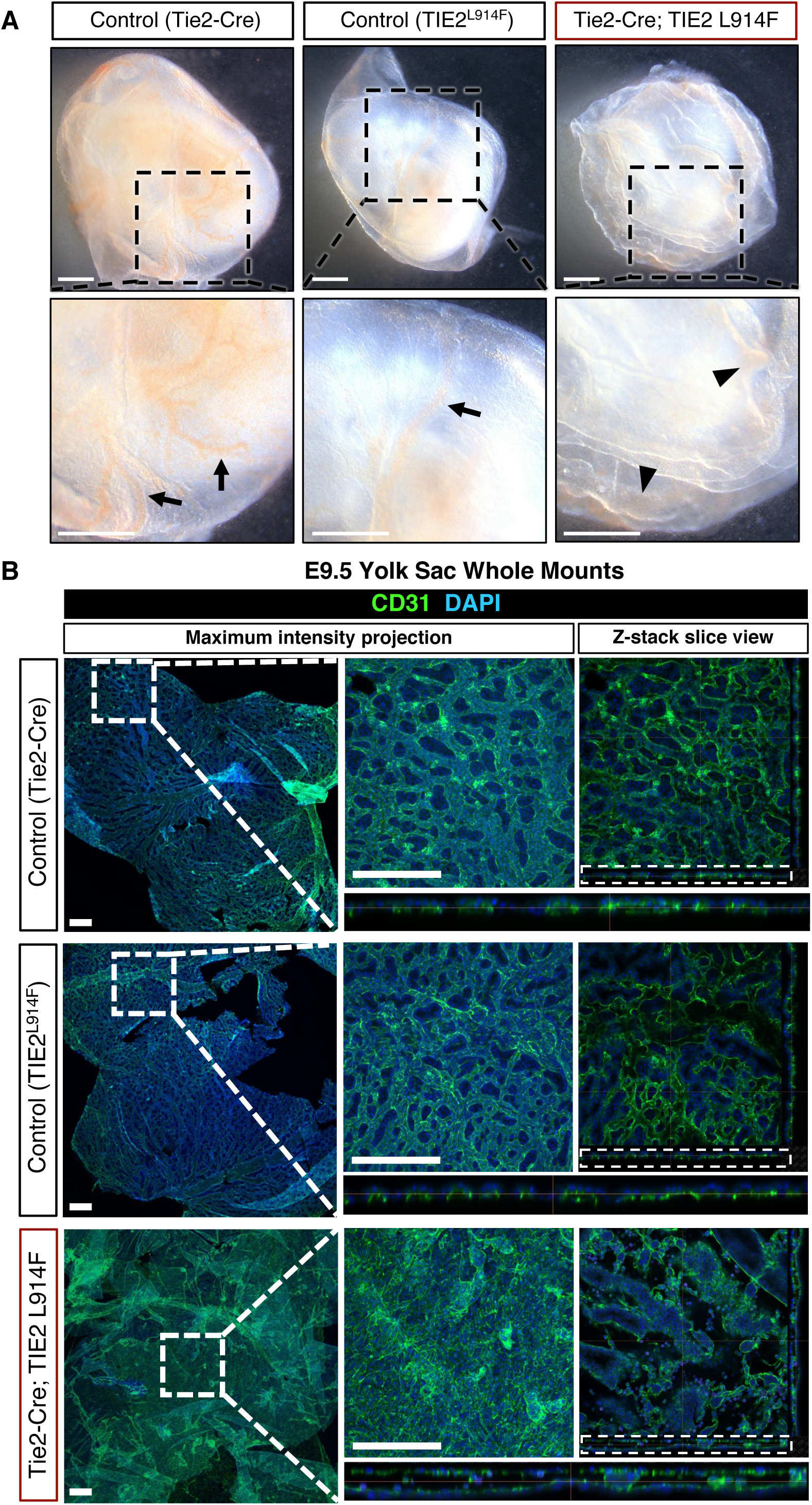
Expression of TIE2^L914F^ interferes with vascular remodeling in E9.5 yolk sacs. (A) Gross stereomicroscope images of embryos and yolk sacs collected at E9.5. Zoomed images in black boxes show the presence of visible large, blood-filled vessels in controls (arrows) and areas of blood pooling in mutants (arrowheads). Scale bars=1mm. (B) Whole mount yolk sacs were immunostained for the vasculature (CD31, green), counterstained with DAPI (blue), and imaged with confocal microscopy. Maximum intensity projections of z-stack images are shown in left images. Rightmost images show a z-stack slice view of high magnification image. Z-plane sections in white boxes are shown enlarged directly beneath slice view images. Representative of n=13 controls and n=7 mutants. Scale bars=250µm.

As TIE2 p.L914F has been implicated in driving increased cell proliferation *in vitro*^47^, here we sought to determine whether the observed changes to the vasculature were due to increased cellular proliferation in *Tie2-Cre;TIE2^L914F^* yolk sacs. We immunostained whole-mount yolk sacs for phosphorylated histone 3 (phospho-H3), a specific marker of condensed chromatin during mitosis^48^. We found no difference in the number of proliferating cells between control and mutant yolk sacs (**Figure 5A** and **B**), indicating that the vessel expansion seen in *Tie2-Cre;TIE2^L914F^*yolk sacs is likely due to errors in EC remodeling and migration, rather than increased proliferation.

**Figure 5:**
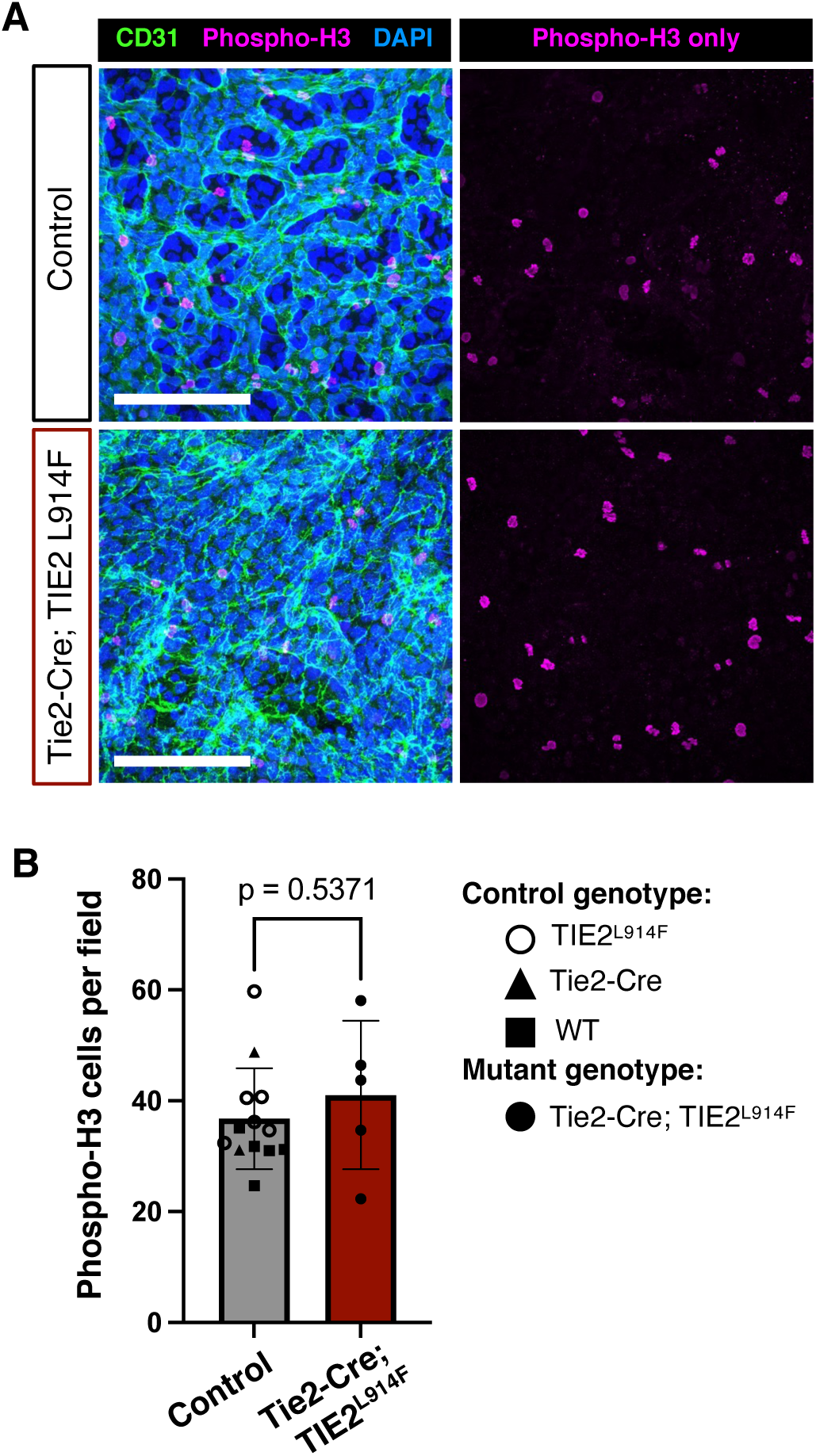
Vascular remodeling defects in E9.5 Tie2-Cre;TIE2^L914F^ yolk sacs are not associated with increased cellular proliferation. (A) E9.5 yolk sacs were immunostained for phospho-histone 3 (phospho-H3), a marker of proliferating cells (magenta), CD31 (blood vessels, green), and DAPI (nuclei, blue). Scale bar=500µm. (B) The number of phospho-H3 cells per image field was quantified. Each data point is the average of 10 images per mouse yolk sac. n=13 controls, n=5 mutants. Control specimens include yolk sacs of each genotype, indicated in the figure legend. Mean±SD. Welch’s t-test.

In total, these data demonstrate that, in the embryonic yolk sac, expression of the mutant TIE2 drives dilation and disorganization of the vascular plexus, without increasing cell proliferation, likely due to impaired vascular remodeling.

### *Tie2-Cre;TIE2^L914F^* yolk sacs retain the ability to produce erythroid cells

The yolk sac is also the site where the first waves of primitive and definitive hematopoiesis take place. The production of hematopoietic precursors is closely tied to the development of the endothelium within the primitive blood islands, with both lineages thought to arise from a common precursor^46,49,50^. Likely due to this shared lineage, *Tie2-Cre* expression has also been reported in hematopoietic cells of the embryo, in addition to its expression within ECs^51^. As defects in hematopoiesis and erythroid cell production have been associated with embryonic lethality at E9.5-11.5^52–55^, we sought to determine whether erythroid cells are present in *Tie2-Cre;TIE2^L914F^* yolk sacs. Upon visual inspection, mutant yolk sacs appeared to contain blood cells, although these were abnormally pooled randomly throughout the yolk sac due to the lack of an organized vascular network (**Figure 4A**).

To confirm the presence of erythroid cells, we performed immunostaining for Ter119, an erythrocyte-specific marker that is expressed early during erythropoiesis^56^. In E9.5 control whole mount yolk sacs, we observed abundant Ter119^+^ cells within the vascular network (**Figure 6A, left**). In *Tie2-Cre;TIE2^L914F^* yolk sacs, Ter119^+^ erythroid cells were still present, but were randomly distributed throughout the abnormally enlarged vascular lumens (**Figure 6A, right**). We further stained and examined the embryos for Ter119 expressing cells. In control embryos we observed many Ter119^+^ cells distributed throughout the vascular system (**Figure 6B, left**). Conversely, in mutant embryos there was a decrease in the number of erythroid cells, likely due to impaired circulation from the yolk sac, which is the primary contributor of hematopoietic cells at this timepoint (**Figure 6B, right**). While there was spatial disorganization and reduced number of Ter119^+^ erythroid cells in *Tie2-Cre;TIE2^L914F^* animals, the presence of these cells at all likely suggest that the initial phases of hematopoiesis are functional.

**Figure 6:**
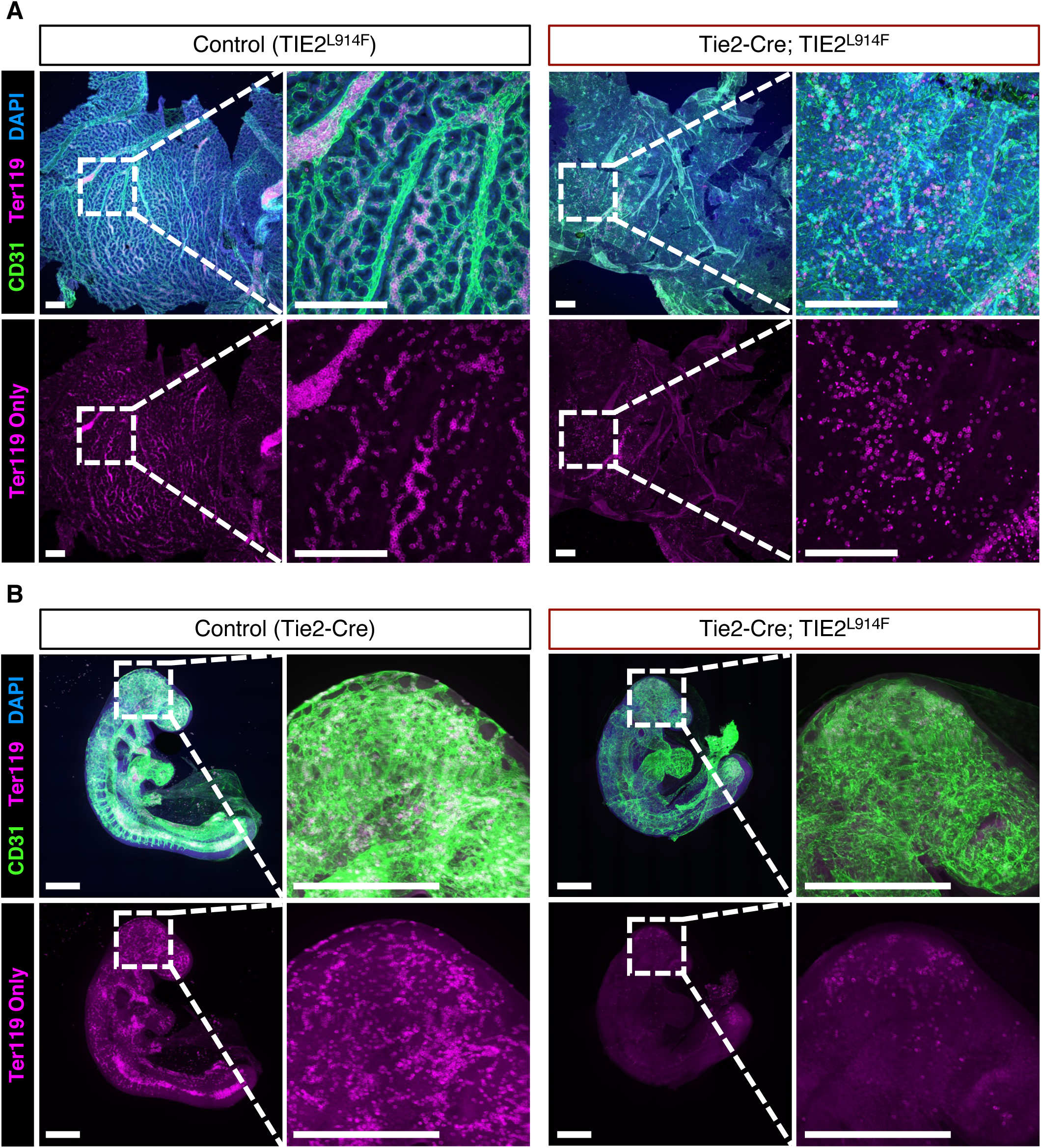
Erythroid cells in Tie2-Cre; TIE2^L914F^ E9.5 yolk sacs and embryos. (A) Whole mount yolk sacs were immunostained for the vasculature (CD31, green), erythroid cells (Ter119, magenta), and nuclei counterstained with DAPI (blue). Maximum intensity projections of z-stack images are shown. White boxes show locations of magnified areas to the right. Representative of n=10 controls and n=4 mutants. Scale bars=250µm. (B) Whole mount embryos were immunostained for CD31, Ter119, and counterstained with DAPI. Images to the left are representative maximum intensity projections of tile-scan z-stack confocal images. Magnified images (white boxes) illustrate Ter119^+^ cells of the cranial region. Scale bars=500µm. Representative of n=7 controls and n=5 mutants.

Together, these data rule out erythroid cell deficiency as the primary driver of embryonic lethality.

## Discussion

Here we generated a novel mouse model that enabled us to experimentally test the hypothesis that constitutive expression of mutant TIE2^*L914F*^ in the developing vasculature causes embryonic lethality. We show that *Tie2-Cre; TIE2^L914F^* embryos fail to develop beyond E9.5, exhibiting malformations of the blood vessels in the embryo and severe remodeling defects in the vasculature of the yolk sac. We further show that the yolk sac remodeling defects are not associated with increased cell proliferation or a total loss of erythroid cells. These experimental data corroborate observations of the genetics of VM patients, where the TIE2 p.L914F mutation is never found in germline, inherited forms of disease due to incompatibility with embryonic survival.

These data also add to the field’s growing body of knowledge on the role of TIE2 signaling in vascular development. Interestingly, the initial stages of vasculogenesis (in which ECs are generated by angioblast precursors and establish de-novo the first vessels of the embryo) appear unaffected by the mutation, even though TIE2 has been reported to be expressed in angioblasts^57^. This is in line with the phenotypes observed in TIE2 knockout mouse studies^30,31^, where loss of TIE2 does not interfere with the establishment of the EC lineage and does not affect the initial stages of the vascular development up to E8.5. Similar to loss of TIE2, we show that expression of hyperactive TIE2 also causes abnormal vascular remodeling and angiogenic defects. Interestingly, in Sato et al., the reported phenotypes are very similar to what was observed in our model. They found that *Tek*-knockout animals have dilation of the vascular networks in the head region as well as in the yolk sac, with the loss of vascular patterning into a hierarchical tree of larger and smaller vessels^31^. Together with the data reported in our study, these seem to indicate: 1) although TIE2 expression is reported in angioblasts, the effects of neither TIE2 deletion nor hyperactivation is sufficient to interfere with vasculogenesis and 2) the balance of TIE2 activation is of utmost importance in vascular development, with both increased and decreased signaling contributing to similar defects in remodeling of the vascular system.

While vasculogenesis in *Tie2-Cre;TIE2^L914F^* mice did occur, producing the rudimentary vasculature of the embryo and yolk sac, the subsequent remodeling of these vessels was significantly impaired by the mutation. Changes to the phenotype of the yolk sac vasculature were readily apparent, with normal remodeling into a patterned vascular tree seeming to be completely inhibited. Instead of forming hierarchical vascular structures, capable of circulating blood to and from the embryo, the ECs in *Tie2-Cre;TIE2^L914F^* mice formed semi-continuous sheets. Similar yolk sac vascular phenotypes have been associated with increased cellular proliferation^21^, however we did not observe this in our model. While the vascular organization was highly abnormal, it is interesting to note that several important characteristics of a normal blood vessel were still present. The layer of ECs was continuous, with robust expression of CD31 at the cell borders. The vessel structures also seemed to be lumenized, with a blood-filled space present between the two sheets of cells. Therefore, while remodeling of the primitive capillary plexus was impeded by expression of the TIE2 p.L914F mutation, the ECs still retained the ability to form lumenized, vessel-like structures and maintain EC-to-EC connections.

Expression of TIE2 p.L914F in embryonic ECs also led to changes to the embryonic and extraembryonic vasculature that are consistent with the role of the mutation in driving sporadic vascular malformation. The embryonic vascular anomalies include improper organization and dilation of the cranial plexus and aberrant expansion of the pharyngeal arch arteries and dorsal aorta. The disorganization of the cranial plexus likely reflects similar remodeling defects as seen in the yolk sac vasculature, where formation of a hierarchical network of large and small vessels from an immature plexus was inhibited. *In-vitro* studies utilizing 3-dimensional lumen formation assays have shown that ECs expressing TIE2^*L914F*^ form structures with increased lumen size^58^, supporting the possibility that the mutant ECs are directly responsible for the increase in lumen diameter of these vessels. In this study, they also showed that *in-vitro* EC migration is altered by the mutation, which may be related to the remodeling defects observed *in-vivo*.

It is interesting to note the dilation of the large vessels, including the pharyngeal arch arteries and dorsal aorta. In the context of inherited and sporadic vascular malformation, TIE2 mutations have only ever been discovered in VM. The ability of TIE2^*L914F*^ to cause malformations of the arterial vessels of the embryo could reflect the active state of these growing vessels, which contrasts with their relatively quiescent state in postnatal and adult tissues^59^, or the fact that vessel-type identity can be more plastic during development^60^. However, it is also well-established that blood flow plays a critical role in control of vessel size during development^40^. As blood flow from the yolk sac would be severely inhibited due to the remodeling defects, it is possible that this change in flow through the large vessels could be responsible for the increase in size. It is therefore difficult to experimentally ascertain whether the arterial defects are due to cell autonomous or non-cell autonomous effects of the mutation. However, there is some level of circulation, as the hearts of mutant embryos still contract and some erythroid cells are present in yolk sacs and embryos, ruling out major cardiac or hematopoietic deficiency as being totally responsible for the vascular defects.

In addition, the phenotypes we observed in our hyperactive TIE2 model recapitulate key aspects of phenotypes observed with other mutations related to vascular malformations and the TIE2 signaling pathway. Mouse embryos deficient in *Foxo1* exhibit similar growth retardation, vascular remodeling defects, and embryonic lethality as observed in our model^61^. Expression of hyperactivating, oncogenic *Pik3ca*^H1047R^ in mouse ECs also presented with embryonic and extraembryonic defects in vascular remodeling, with associated embryonic death^21^. Furthermore, mouse embryos deficient in VE-PTP, a phosphatase targeting TIE2^62^ , exhibit strikingly similar defects in yolk sac vasculature^63^. WT and mutant TIE2 are known to signal through the PI3K/AKT/mTOR axis^34,64,65^, which causes downstream phosphorylation and inactivation of FOXO1^15,66^. Furthermore, this pathway is also of great interest due to its associations to the formation of vascular malformations, especially VM. TIE2 hyperactivating mutations were the first to be associated with VM and have since been associated with a large percentage of VM cases^8–11^. However, more recently, hyperactivating mutations in *PIK3CA* have also been identified in VM patients^67,68^ and expression of mutant *PIK3CA* expression in blood ECs has successfully recapitulated the VM phenotype in mouse models^69,70^. The similarities in these phenotypes (both developmental and pathological) and the fact that these mutations appear to converge on a shared pathway suggest perhaps a common cellular mechanism in these models and phenotypes. This also emphasizes the importance of this pathway in regulating normal vascular development and remodeling.

There are some limitations to the model used in these studies. Firstly, the *TIE2^L914F^*mouse line utilizes a transgene which is controlled by the *Rosa26* promoter, rather than an endogenous *Tek* promoter. This may result in differences in the transcriptional regulation of the transgene. However, in this model, we utilized *Tie2-Cre* to control activation of the transgene, which does utilize a *Tek* promoter region, restoring some dependence of the transgene on normal *Tek* transcriptional timing during development. Secondly, as the embryonic vascular phenotype is severe and ultimately results in mid-gestation embryonic death, it is difficult to distinguish between cell-autonomous and non-cell autonomous effects of the mutation. Based on the phenotypes reported in the literature utilizing *Tek* knockout mice and mice with hyperactivating mutations in related proteins (see discussion points above), we can confidently assign the vascular remodeling defects to the direct action of the TIE2 mutation in endothelial cells. However, other phenotypes, including the dilation of the large vessels (dorsal aorta and pharyngeal arch arteries) could possibly be due to insufficient circulation stemming from the yolk sac vascular defects and growth retardation preceding embryonic demise. Future studies utilizing tools to induce mosaic transgene expression could answer these questions.

In conclusion, the mouse model developed in this study represents a novel genetic animal model of VM-associated TIE2 hyperactivating mutation. We demonstrate that early developmental expression of TIE2^*L914F*^ is embryonically lethal, providing experimental evidence to explain why this mutation is only associated with sporadic, and never hereditary, disease. We found that the mutation interferes with vascular remodeling, contributing to the field’s knowledge of the role of TIE2 in this process.

In conclusion, we foresee that this model will be an important tool for future studies aimed at dissecting TIE2 signaling, VM pathogenesis, and to perform pre-clinical testing of therapeutics for VM patients.

## Experimental procedures

### Institutional permissions

All mouse experiments and procedures have been reviewed and approved by the CCHMC Institutional Animal Care and Use Committee (protocol number IACUC 2023-0025). All animals were cared for in accordance with guidelines from the National Institutes of Health. Studies were performed using both male and female mice.

### Generation and genotyping of B6-Tg(Rosa26-TIE2^L914F^)EBos mice

A targeting vector was constructed that contains the minimal adenovirus type 2 major late splice acceptor, a Neo-stop cassette flanked by *loxP* sites (floxed), the coding sequence of the human *TEK* gene with the p.L914F mutation, a woodchuck hepatitis virus posttranscriptional regulatory element (WPRE), and a bovine growth hormone polyadenylation signal (Ingenious Targeting Laboratory). The *TEK^L914F^* gene is followed by an IRES sequence and mCherry fluorescent reporter, which allows for mCherry to be co-expressed with mutant TIE2 using the same promoter. The entire construct was flanked by short and long *Rosa26* homology arms for targeted integration into the *Rosa26* locus.

The transgenic construct was transfected into iTL IN2 (C57BL/6) embryonic stem cells. Positive clones were then microinjected into Balb/c blastocysts. Chimeras with high percentage black coat color were mated with wild-type C57BL/6N mice to generate F1 heterozygous offspring. PCR for various components of the construct was used to identify pups with successful integration of the transgene.

For routine genotyping of colony and experimental *TIE2^L914F^* mice, a three-primer PCR protocol was developed to simultaneously amplify the wild-type *Rosa26* locus (spanning the integration site) and the 5’ end of the transgenic construct. The primers utilized were: Rosa_SQ1 (AGC ACT TGC TCT CCC AAA GTC), Rosa_WT R (AAT CTG TGG GAA GTC TTG TCC), and LAN1 (CCA GAG GCC ACT TGT GTA GC). Each PCR reaction contained 1µM Rosa_SQ1, 0.5µM Rosa_WT R, 0.5µM LAN1, 1x EconoTaq Plus green 2X Master Mix (Lucigen), and 2µl mouse DNA in 15µl total volume. The reaction parameters consist of 94°C (2min), 40 cycles of 94°C (30s), 64°C (30s), 72°C (1min), followed by one hold at 72°C (2min). The wild-type *Rosa26* allele amplifies as a ∼330bp product and the mutant transgene amplifies as a ∼500bp product, when run with agarose gel electrophoresis. For colony genotyping, murine DNA was extracted from ear biopsies using the protocol established in Truett, et al. (2000)^71^. For genotyping of mouse embryos, either the yolk sac or the embryo tail was processed using the same protocol.

### Mouse husbandry and embryo collection

To generate experimental offspring, female *B6-Tg(Rosa26-TIE2^L^*^914^*^F^)^EBos^* mice were crossed with male *B6.Cg-Tg(Tek-cre)1^Ywa/J^* mice, described previously in Kisanuki, et al. (2001)^36^. Mendelian ratios were calculated from offspring genotypes at weaning.

For embryo studies, timed breeding was performed. Female mice were group-housed and exposed to male bedding three nights before mating to synchronize estrous cycles (Whitten effect^72^). Upon combining mating pairs, females were examined each morning for the presence of a vaginal plug. The day that the plug was found was considered E0.5.

Pregnant females were euthanized before removal of embryo implantation sites, which were subsequently stored in HBSS. Embryos and yolk sacs were dissected and imaged using a Leica S8 APO stereomicroscope equipped with a Flexacam C3 camera. Crown-rump measurements were obtained using ImageJ software.

E9.5 embryos and yolk sacs were fixed in 10% neutral-buffered formalin for 1h at room temperature and subsequently stored in PBS at 4°C. Older embryos (E11.5 and 13.5) were fixed overnight at 4°C.

### Sanger sequencing of mouse and human *TEK*

Genotyping DNA samples were used for amplification of Exon 17 of both the endogenous mouse *Tek* gene and the human *TEK* transgene. The mouse gene was amplified with the following primers: Forward (GAT CCC TGC TAG GCA GTT GT) and Reverse (CTC AAG TAG TCC ATC CCC CG). The following forward primer was used for Sanger sequencing of the region containing leucine 913: AGG GAC TCT GAT GAA GGT AAG AAG A. The human gene was amplified with the following primers: Forward (TGG GTT ACG GAT GGA TGC TG) and Reverse (ACC CCT AGG AAT GCT CGT TCA). The forward primer was also used for Sanger sequencing of the region containing leucine 914. The PCR reaction was carried out according to manufacturer’s instruction using Q5 Hot Start High-Fidelity 2X Master Mix (NEB #M0494). Sanger sequencing was performed by the CCHMC Genomics Sequencing Facility (RRiD SCR_022630) Chromatograms were visualized using SnapGene software (www.snapgene.com).

### Whole-mount immunostaining of embryos and yolk sacs

For immunostaining, 5% BSA / 0.5% Triton-X in PBS was used as blocking and antibody buffers. After fixation, whole embryos and yolk sacs were washed in PBS and incubated in blocking buffer for 1h at room temperature. Tissues were then incubated sequentially in primary and secondary antibodies, each overnight at 4°C with PBS washes in between. The following primary antibodies were used: CD31/PECAM1 Alexa Fluor® 488-conjugated (R&D #FAB3628, 1:100), phospho-Histone 3 (Millipore #06-570, 1:100), TER-119 Alexa Fluor® 647-conjugated (Biolegend #116218, 1:100). For phospho-Histone 3 staining, the following secondary antibody was used: Goat anti-rabbit IgG Alexa Fluor® Plus 594 (Invitrogen #A32740, 1:200). Nuclei counterstaining was performed with DAPI (4’,6-diamidino-2-phenylindole) (Invitrogen #D1306, 5µg/mL). Yolk sac samples were mounted on slides using Fluoromount-G® mounting medium (Southern Biotech, #0100-01) and sealed with a coverslip. Embryo samples were cleared for 1h in EZ View reagent^73^ and then mounted on slides in the same solution. To hold the embryos, wells were created with Secure-Seal™ spacers (Invitrogen, #S24737), which were stacked using at least five layers to achieve a 500µm well space.

### Confocal imaging and quantification

Z-stack images of embryos and yolk sacs were acquired on a Nikon A1 Inverted LUNV laser-scanning confocal microscope. For phospho-H3 analysis, 10 randomly assigned regions were imaged per sample. Nikon Elements software was used to create maximum intensity projections or depth-coded volume views of the images. Fiji (ImageJ)^74^ software was used to quantify the number of phospho-H3 positive nuclei per field.

### Data collection and statistics

Prism 9.0 software (GraphPad) was used to make graphs and perform statistical analysis. Data are presented as Mean±standard deviation with n values reported in figure legends. Statistical significance between two groups was assessed by unpaired Welch’s t-test. Statistical significance is defined as p values less than 0.05. Schematics were created with Biorender.com

## Acknowledgements and sources of funding

Research reported in this manuscript was supported by the National Heart, Lung, and Blood Institute, under Award Numbers R01HL117952 (E.B.), R01HL167700 (E.B.), and F31HL176101 (L.J.B.), part of the National Institutes of Health. The content is solely the responsibility of the authors and does not necessarily represent the official views of the National Institutes of Health.

Additional funding supporting the study was provided by the American Heart Institute (AHA) Predoctoral Fellowship (https://doi.org/10.58275/AHA.24PRE1191403.pc.gr.190588) to L.J. Bischoff. We thank Dr. Katherine Yutzey for critical review of the manuscript. We thank the Bio-imaging and Analysis Facility and Veterinary Services at Cincinnati Children’s Hospital Medical Center for providing state-of-the-art instrumentation, services, training, and education.

## Author contributions

L.J.B, S.S., and E.B. conceived the project. E.B. supervised the research. L.J.B., S.S., and K.S. performed the mouse experiments and sample collection. L.J.B. performed immunostaining, confocal imaging, and data analysis. L.J.B. wrote the manuscript and prepared the figures. All authors revised and commented on the manuscript.

## References

1. Wassef M, Blei F, Adams D, et al. Vascular Anomalies Classification: Recommendations From the International Society for the Study of Vascular Anomalies. Pediatrics. 2015;136(1):e203–214. doi:10.1542/peds.2014-3673

2. Vikkula M, Boon LM, Mulliken JB. Molecular genetics of vascular malformations. Matrix Biol. 2001;20(5-6):327–335. doi:10.1016/S0945-053X(01)00150-0

3. Behravesh S, Yakes W, Gupta N, et al. Venous malformations: Clinical diagnosis and treatment. Cardiovasc Diagn Ther. 2016;6(6):557–569. doi:10.21037/cdt.2016.11.10

4. Vogel SA, Hess CP, Dowd CF, et al. Early versus later presentations of venous malformations: Where and why? Pediatr Dermatol. 2013;30(5):534–540. doi:10.1111/pde.12162

5. Dompmartin A, Vikkula M, Boon LM. Venous Malformation: update on etiopathogenesis, diagnosis & management. Phlebol Venous Forum R Soc Med. 2010;25(5):224. doi:10.1258/PHLEB.2009.009041

6. Swerdlin RF, Briones MA, Gill AE, Hawkins CM. Coagulopathy and related complications following sclerotherapy of congenital venous malformations. Pediatr Blood Cancer. 2022;69(5):e29610. doi:10.1002/pbc.29610

7. Dompmartin A, Acher A, Thibon P, et al. Association of localized intravascular coagulopathy with venous malformations. Arch Dermatol. 2008;144(7):873–877. doi:10.1001/archderm.144.7.873

8. Vikkula M, Boon LM, Carraway KL, et al. Vascular Dysmorphogenesis Caused by an Activating Mutation in the Receptor Tyrosine Kinase TIE2. Cell. 1996;87:1181–1190.

9. Soblet J, Limaye N, Uebelhoer M, Boon LM, Vikkula M. Variable somatic TIE2 mutations in half of sporadic venous malformations. Mol Syndromol. 2013;4(4):179–183. doi:10.1159/000348327

10. Calvert JT, Riney TJ, Kontos CD, et al. Allelic and Locus Heterogeneity in Inherited Venous Malformations. Hum Mol Genet. 1999;8(7):1279–1289. doi:10.1093/hmg/8.7.1279

11. Limaye N, Wouters V, Uebelhoer M, et al. Somatic Mutations in the Angiopoietin-Receptor TIE2 Can Cause Both Solitary and Multiple Sporadic Venous Malformations. Nat Genet. 2009;41(1):118–124. doi:10.1038/ng.272

12. Borst AJ, Nakano TA, Blei F, Adams DM, Duis J. A Primer on a Comprehensive Genetic Approach to Vascular Anomalies. Front Pediatr. 2020;8. doi:10.3389/fped.2020.579591

13. Goines J, Li X, Cai Y, et al. A xenograft model for venous malformation. Angiogenesis. 2018;21(4):725–735. doi:10.1007/s10456-018-9624-7

14. Wouters V, Limaye N, Uebelhoer M, et al. Hereditary cutaneomucosal venous malformations are caused by TIE2 mutations with widely variable hyper-phosphorylating effects. Eur J Hum Genet. 2010;18(4):414–420. doi:10.1038/ejhg.2009.193

15. Uebelhoer M, Nätynki M, Kangas J, et al. Venous malformation-causative TIE2 mutations mediate an AKT-dependent decrease in PDGFB. Hum Mol Genet. 2013;22(17):3438–3448. doi:10.1093/hmg/ddt198

16. Nätynki M, Kangas J, Miinalainen I, et al. Common and specific effects of TIE2 mutations causing venous malformations. Hum Mol Genet. 2015;24(22):6374. doi:10.1093/HMG/DDV349

17. Happle R. Lethal genes surviving by mosaicism: A possible explanation for sporadic birth defects involving the skin. J Am Acad Dermatol. 1987;16(4):899–906. doi:10.1016/S0190-9622(87)80249-9

18. Martínez-Glez V, Tenorio J, Nevado J, et al. A Six-attribute Classification of Genetic Mosaicism. Genet Med Off J Am Coll Med Genet. 2020;22(11):1743–1757. doi:10.1038/s41436-020-0877-3

19. Wetzel-Strong SE, Galeffi F, Benavides C, et al. Developmental expression of the Sturge–Weber syndrome-associated genetic mutation in Gnaq: a formal test of Happle’s paradominant inheritance hypothesis. Genetics. 2023;224(4):iyad077. doi:10.1093/genetics/iyad077

20. Schrenk S, Bischoff LJ, Goines J, et al. MEK inhibition reduced vascular tumor growth and coagulopathy in a mouse model with hyperactive GNAQ. Nat Commun. 2023;14(1):1929. doi:10.1038/s41467-023-37516-7

21. Hare LM, Schwarz Q, Wiszniak S, et al. Heterozygous expression of the oncogenic *Pik3ca*H1047R mutation during murine development results in fatal embryonic and extraembryonic defects. Dev Biol. 2015;404(1):14–26. doi:10.1016/j.ydbio.2015.04.022

22. Schnürch H, Risau W. Expression of tie-2, a member of a novel family of receptor tyrosinekinases, in the endothelial cell lineage. Development. 1993;119(3):957–968.

23. Eklund L, Kangas J, Saharinen P. Angiopoietin–Tie signalling in the cardiovascular and lymphatic systems. Clin Sci Lond Engl 1979. 2017;131(1):87-103. doi:10.1042/CS20160129

24. Fukuhara S, Sako K, Noda K, Nagao K, Miura K, Mochizuki N. Tie2 is tied at the cell-cell contacts and to extracellular matrix by Angiopoietin-1. Exp Mol Med. 2009;41(3):133–139. doi:10.3858/emm.2009.41.3.016

25. Kim M, Allen B, Korhonen EA, et al. Opposing actions of angiopoietin-2 on Tie2 signaling and FOXO1 activation. J Clin Invest. 2016;126(9):3511–3525. doi:10.1172/JCI84871

26. Savant S, La Porta S, Budnik A, et al. The Orphan Receptor Tie1 Controls Angiogenesis and Vascular Remodeling by Differentially Regulating Tie2 in Tip and Stalk Cells. Cell Rep. 2015;12(11):1761–1773. doi:10.1016/j.celrep.2015.08.024

27. Wong AL, Haroon ZA, Werner S, Dewhirst MW, Greenberg CS, Peters KG. Tie2 Expression and Phosphorylation in Angiogenic and Quiescent Adult Tissues. Circ Res. 1997;81(4):567–574. doi:10.1161/01.RES.81.4.567

28. Bogdanovic E, Nguyen VPKH, Dumont DJ. Activation of Tie2 by angiopoietin-1 and angiopoietin-2 results in their release and receptor internalization. J Cell Sci. 2006;119(17):3551–3560. doi:10.1242/jcs.03077

29. Seegar TCM, Eller B, Tzvetkova-Robev D, et al. Tie1-Tie2 Interactions Mediate Functional Differences between Angiopoietin Ligands. Mol Cell. 2010;37(5):643–655. doi:10.1016/j.molcel.2010.02.007

30. Dumont DJ, Gradwohl G, Fong GH, et al. Dominant-negative and targeted null mutations in the endothelial receptor tyrosine kinase, tek, reveal a critical role in vasculogenesis of the embryo. Genes Dev. 1994;8(16):1897–1909. doi:10.1101/gad.8.16.1897

31. Sato TN, Tozawa Y, Deutsch U, et al. Distinct roles of the receptor tyrosine kinases Tie-1 and Tie-2 in blood vessel formation. Nature. 1995;376(6535):70-74. doi:10.1038/376070a0

32. Suri C, Jones PF, Patan S, et al. Requisite Role of Angiopoietin-1, a Ligand for the TIE2 Receptor, during Embryonic Angiogenesis. Cell. 1996;87(7):1171–1180. doi:10.1016/S0092-8674(00)81813-9

33. Maisonpierre PC, Suri C, Jones PF, et al. Angiopoietin-2, a Natural Antagonist for Tie2 That Disrupts in vivo Angiogenesis. Science. 1997;277(5322):55-60. doi:10.1126/science.277.5322.55

34. Boscolo E, Limaye N, Huang L, et al. Rapamycin improves TIE2-mutated venous malformation in murine model and human subjects. J Clin Invest. 2015;125(9):3491. doi:10.1172/JCI76004

35. Goines J, Boscolo E. A Xenograft Model for Venous Malformation. Methods Mol Biol Clifton NJ. 2021;2206:179–192. doi:10.1007/978-1-0716-0916-3_13

36. Kisanuki YY, Hammer RE, Miyazaki J, Williams SC, Richardson JA, Yanagisawa M. Tie2-Cre transgenic mice: a new model for endothelial cell-lineage analysis in vivo. Dev Biol. 2001;230(2):230–242. doi:10.1006/dbio.2000.0106

37. Walls JR, Coultas L, Rossant J, Henkelman RM. Three-Dimensional Analysis of Vascular Development in the Mouse Embryo. PLoS ONE. 2008;3(8):e2853. doi:10.1371/journal.pone.0002853

38. Hiruma T, Nakajima Y, Nakamura H. Development of pharyngeal arch arteries in early mouse embryo. J Anat. 2002;201(1):15–29. doi:10.1046/j.1469-7580.2002.00071.x

39. Lucitti JL, Jones EAV, Huang C, Chen J, Fraser SE, Dickinson ME. Vascular remodeling of the mouse yolk sac requires hemodynamic force. Dev Camb Engl. 2007;134(18):3317–3326. doi:10.1242/dev.02883

40. Udan RS, Vadakkan TJ, Dickinson ME. Dynamic responses of endothelial cells to changes in blood flow during vascular remodeling of the mouse yolk sac. Dev Camb Engl. 2013;140(19):4041–4050. doi:10.1242/dev.096255

41. Huang C, Sheikh F, Hollander M, et al. Embryonic atrial function is essential for mouse embryogenesis, cardiac morphogenesis and angiogenesis. Development. 2003;130(24):6111–6119. doi:10.1242/dev.00831

42. May SR, Stewart NJ, Chang W, Peterson AS. A *Titin* mutation defines roles for circulation in endothelial morphogenesis. Dev Biol. 2004;270(1):31–46. doi:10.1016/j.ydbio.2004.02.006

43. de Boer BA, van den Berg G, de Boer PAJ, Moorman AFM, Ruijter JM. Growth of the developing mouse heart: An interactive qualitative and quantitative 3D atlas. Dev Biol. 2012;368(2):203–213. doi:10.1016/j.ydbio.2012.05.001

44. Eichmann A, Corbel C, Nataf V, Vaigot P, Bréant C, Le Douarin NM. Ligand-dependent development of the endothelial and hemopoietic lineages from embryonic mesodermal cells expressing vascular endothelial growth factor receptor 2. Proc Natl Acad Sci. 1997;94(10):5141–5146. doi:10.1073/pnas.94.10.5141

45. Ferkowicz MJ, Starr M, Xie X, et al. CD41 expression defines the onset of primitive and definitive hematopoiesis in the murine embryo. Development. 2003;130(18):4393–4403. doi:10.1242/dev.00632

46. Haar JL, Ackerman GA. A phase and electron microscopic study of vasculogenesis and erythropoiesis in the yolk sac of the mouse. Anat Rec. 1971;170(2):199–223. doi:10.1002/ar.1091700206

47. Li X, Cai Y, Goines J, et al. Ponatinib Combined With Rapamycin Causes Regression of Murine Venous Malformation. Arterioscler Thromb Vasc Biol. 2019;39(3):496. doi:10.1161/ATVBAHA.118.312315

48. Dai J, Sultan S, Taylor SS, Higgins JMG. The kinase haspin is required for mitotic histone H3 Thr 3 phosphorylation and normal metaphase chromosome alignment. Genes Dev. 2005;19(4):472–488. doi:10.1101/gad.1267105

49. Gonzalez-Crussi F. Vasculogenesis in the chick embryo. An ultrastructural study. Am J Anat. 1971;130(4):441–459. doi:10.1002/aja.1001300406

50. Shalaby F, Ho J, Stanford WL, et al. A Requirement for Flk1 in Primitive and Definitive Hematopoiesis and Vasculogenesis. Cell. 1997;89(6):981–990. doi:10.1016/S0092-8674(00)80283-4

51. Tang Y, Harrington A, Yang X, Friesel RE, Liaw L. The contribution of the Tie2+ lineage to primitive and definitive hematopoietic cells. Genes N Y N 2000. 2010;48(9):563-567. doi:10.1002/dvg.20654

52. Tsang AP, Fujiwara Y, Hom DB, Orkin SH. Failure of megakaryopoiesis and arrested erythropoiesis in mice lacking the GATA-1 transcriptional cofactor FOG. Genes Dev. 1998;12(8):1176–1188. doi:10.1101/gad.12.8.1176

53. Robb L, Lyons I, Li R, et al. Absence of yolk sac hematopoiesis from mice with a targeted disruption of the scl gene. Proc Natl Acad Sci U S A. 1995;92(15):7075–7079.

54. Fujiwara Y, Browne CP, Cunniff K, Goff SC, Orkin SH. Arrested development of embryonic red cell precursors in mouse embryos lacking transcription factor GATA-1. Proc Natl Acad Sci U S A. 1996;93(22):12355–12358.

55. Tsai FY, Keller G, Kuo FC, et al. An early haematopoietic defect in mice lacking the transcription factor GATA-2. Nature. 1994;371(6494):221-226. doi:10.1038/371221a0

56. Socolovsky M, Nam H song, Fleming MD, Haase VH, Brugnara C, Lodish HF. Ineffective erythropoiesis in Stat5a−/−5b−/− mice due to decreased survival of early erythroblasts. Blood. 2001;98(12):3261–3273. doi:10.1182/blood.V98.12.3261

57. Furuta C, Ema H, Takayanagi S ichiro, et al. Discordant developmental waves of angioblasts and hemangioblasts in the early gastrulating mouse embryo. Development. 2006;133(14):2771–2779. doi:10.1242/dev.02440

58. Cai Y, Schrenk S, Goines J, Davis GE, Boscolo E. Constitutive Active Mutant TIE2 Induces Enlarged Vascular Lumen Formation with Loss of Apico-basal Polarity and Pericyte Recruitment. Sci Rep. 2019;9:12352. doi:10.1038/s41598-019-48854-2

59. Lee HW, Xu Y, He L, et al. Role of Venous Endothelial Cells in Developmental and Pathologic Angiogenesis. Circulation. 2021;144(16):1308–1322. doi:10.1161/CIRCULATIONAHA.121.054071

60. Fish JE, Wythe JD. The molecular regulation of arteriovenous specification and maintenance. Dev Dyn. 2015;244(3):391–409. doi:10.1002/dvdy.24252

61. Furuyama T, Kitayama K, Shimoda Y, et al. Abnormal Angiogenesis in Foxo1 (Fkhr)-deficient Mice *. J Biol Chem. 2004;279(33):34741–34749. doi:10.1074/jbc.M314214200

62. Fachinger G, Deutsch U, Risau W. Functional interaction of vascular endothelial-protein-tyrosine phosphatase with the Angiopoietin receptor Tie-2. Oncogene. 1999;18(43):5948–5953. doi:10.1038/sj.onc.1202992

63. Dominguez MG, Hughes VC, Pan L, et al. Vascular endothelial tyrosine phosphatase (VE-PTP)-null mice undergo vasculogenesis but die embryonically because of defects in angiogenesis. Proc Natl Acad Sci. 2007;104(9):3243–3248. doi:10.1073/pnas.0611510104

64. Kontos CD, Stauffer TP, Yang WP, et al. Tyrosine 1101 of Tie2 Is the Major Site of Association of p85 and Is Required for Activation of Phosphatidylinositol 3-Kinase and Akt. Mol Cell Biol. 1998;18(7):4131. doi:10.1128/MCB.18.7.4131

65. Morris PN, Dunmore BJ, Tadros A, et al. Functional analysis of a mutant form of the receptor tyrosine kinase Tie2 causing venous malformations. J Mol Med. 2005;83(1):58–63. doi:10.1007/S00109-004-0601-9/FIGURES/5

66. Daly C, Wong V, Burova E, et al. Angiopoietin-1 modulates endothelial cell function and gene expression via the transcription factor FKHR (FOXO1). Genes Dev. 2004;18(9):1060–1071. doi:10.1101/gad.1189704

67. Limaye N, Kangas J, Mendola A, et al. Somatic Activating PIK3CA Mutations Cause Venous Malformation. Am J Hum Genet. 2015;97(6):914–921. doi:10.1016/j.ajhg.2015.11.011

68. Castel P, Carmona FJ, Grego-Bessa J, et al. Somatic PIK3CA mutations as a driver of sporadic venous malformations. Sci Transl Med. 2016;8(332):332ra42. doi:10.1126/SCITRANSLMED.AAF1164

69. Castillo SD, Tzouanacou E, Zaw-Thin M, et al. Somatic Activating Mutations in Pik3ca Cause Sporadic Venous Malformations in Mice and Humans. Sci Transl Med. 2016;8(332):332ra43. doi:10.1126/scitranslmed.aad9982

70. Petkova M, Kraft M, Stritt S, et al. Immune-interacting lymphatic endothelial subtype at capillary terminals drives lymphatic malformation. J Exp Med. 2023;220(4):e20220741. doi:10.1084/jem.20220741

71. Truett GE, Heeger P, Mynatt RL, Truett AA, Walker JA, Warman ML. Preparation of PCR-Quality Mouse Genomic DNA with Hot Sodium Hydroxide and Tris (HotSHOT). BioTechniques. 2000;29(1):52–54. doi:10.2144/00291bm09

72. Whitten WK. MODIFICATION OF THE OESTROUS CYCLE OF THE MOUSE BY EXTERNAL STIMULI ASSOCIATED WITH THE MALE. Published online July 1, 1956. doi:10.1677/joe.0.0130399

73. Hsu CW, Cerda J III, Kirk JM, et al. EZ Clear for simple, rapid, and robust mouse whole organ clearing. Monk K, Bronner ME, Cox TC, eds. eLife. 2022;11:e77419. doi:10.7554/eLife.77419

74. Schindelin J, Arganda-Carreras I, Frise E, et al. Fiji: an open-source platform for biological-image analysis. Nat Methods. 2012;9(7):676–682. doi:10.1038/nmeth.2019

