## Supplemental data for "Expression of mutant TIE2 p.L914F during mouse development causes embryonic lethality and defects in vascular remodeling"

**Supplemental Figure 1: Optical sectioning of whole-mount Tie2-Cre;TIE2$^{L914F}$ embryos reveals abnormal dilation of the pharyngeal arch arteries and dorsal aorta at E9.5.** The pharyngeal arch areas of the embryos from Figure 3 are shown with volume sections of the YZ plane (blue boxes) and XZ plane (magenta boxes). Yellow lines indicate where the sections are located on the XY plane. Pharyngeal arch arteries are marked with arrowheads and dorsal aortas with arrows.

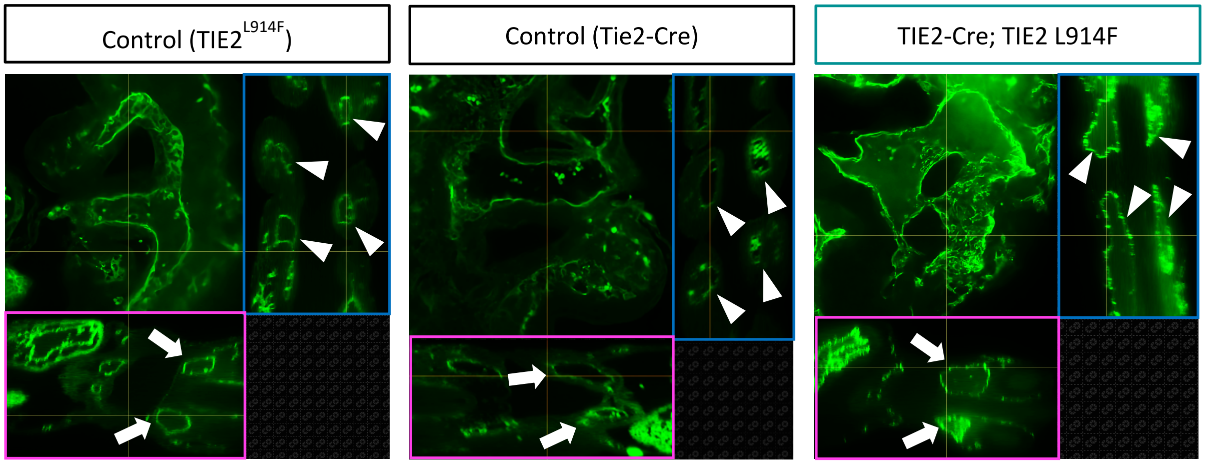

**Supplemental Figure 2: Intersomitic vessels and hearts of Tie2-Cre;TIE2$^{L914F}$ embryos are similar to those in controls at E9.5.** The somites and hearts of the whole-mount embryos from Figure 3 are shown. The vasculature was visualized by CD31 immunostaining. In these volume images, CD31 labeling is depth-coded with rainbow colors according to location on the Z-axis. A and V label the atria and ventricles, respectively.

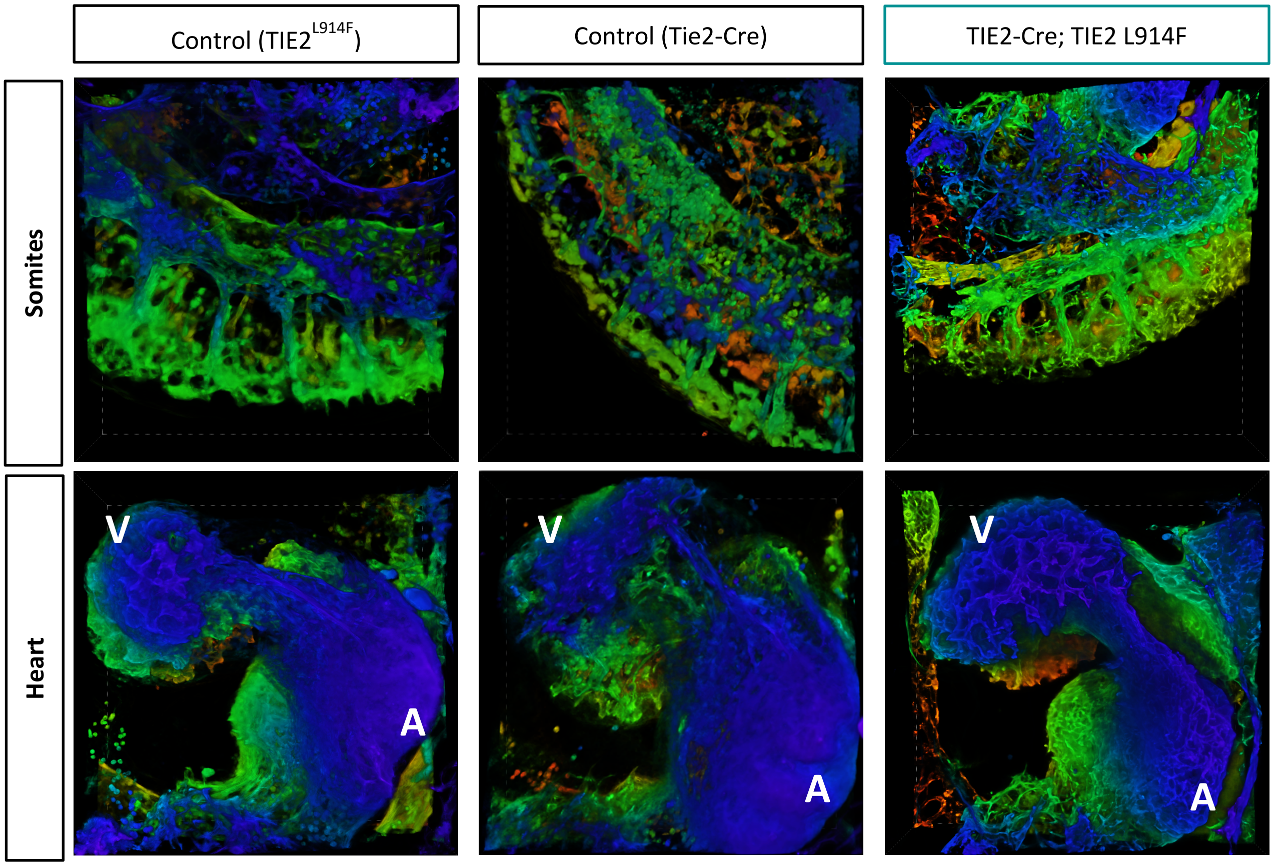
